# Overcoming Fluorescence Loss in mEOS-based AAA+ Unfoldase Reporters Through Covalent Linkage

**DOI:** 10.1101/2025.01.14.633048

**Authors:** Isabella R. Walter, Baylee A. Smith, Dominic Castanzo, Matthew L. Wohlever

**Affiliations:** Department of Chemistry & Biochemistry, University of Toledo, Toledo, OH 43606; Department of Molecular and Cell Biology, University of California, Berkeley, CA 94720; Institute of Quantitative Biosciences, University of California, Berkeley, CA 94720; Department of Cell Biology, University of Pittsburgh, Pittsburgh, PA, 15261

**Keywords:** AAA+ protein, fluorescent protein, proteostasis, membrane protein, protein engineering

## Abstract

Recent work has demonstrated that the soluble photoconvertable fluorescent protein mEOS can be a reporter for AAA+ (ATPases Associated with diverse cellular Activities) unfoldase activity. Given that many AAA+ proteins process membrane proteins, we sought to adapt mEOS for use with membrane protein substrates. However, direct genetic fusion of mEOS to a membrane protein completely abolished fluorescence, severely limiting the utility of mEOS for studying AAA+ proteins. To circumvent this challenge, we separately purified mEOS and a AAA+ degron, covalently linked them via Sortase, and photoconverted the linked construct. This innovative approach preserves fluorescence and enables functional analysis, offering a broadly applicable platform for the study of membrane associated AAA+ proteins.

## Introduction

The functional integrity of the membrane proteins depends on accurate targeting, proper folding, and effective maintenance within the cellular environment[1–3]. Mislocalized or damaged membrane proteins are identified and processed by the cellular proteostasis machinery[4–12]. Defects in this essential process are associated with numerous diseases, including cancer, Parkinson’s Disease, Alzheimer’s disease, and Amyotrophic Lateral Sclerosis[13–16]. The AAA+ (ATPases Associated with diverse cellular Activities) protein family plays a pivotal role in regulating quality control of membrane proteins by assisting in their unfolding, disaggregation, and degradation[17–19]. AAA+ proteins generally form ring shaped hexamers and undergo ATP-dependent conformational changes to translocate substrates through an axial pore, resulting in the mechanical unfolding of the substrate[20,21].

Msp1 is a AAA+ enzyme anchored on the outer mitochondrial membrane (OMM) that helps maintain proteostasis by removing mislocalized proteins from the OMM or substrates that stall in the translocase of the outer membrane (TOM complex)[22–28]. Loss of Msp1 leads to an accumulation of mislocalized proteins on the OMM, mitochondrial fragmentation, and failures in oxidative phosphorylation[29,30]. In addition to the aforementioned quality control roles, the human homolog ATAD1 is also involved in regulating apoptosis via extraction of the pro-apoptotic protein BIM[31]. Developing a robust assay to monitor Msp1/ATAD1 activity is essential for advancing our understanding of their roles in mitochondrial function and human health.

A major challenge in studying Msp1 activity is the lack of a real time assay for Msp1 substrate processing[32]. A common strategy for monitoring substrate processing by AAA+ proteases is to fuse a recognition sequence (degron) to the model substrate GFP because proteolysis leads to a loss of fluorescence[21,33,34]. However, GFP is a poor model substrate for unfoldases such as Msp1 because the lack of a protease domain allows GFP to rapidly refold[35]. Recent work has demonstrated that the soluble, photocovertable fluorescent protein mEOS can be a tool for kinetic analysis of unfoldase activity by AAA+ proteins, including Msp1[36,37]. Photoconversion of mEOS results in a green to red color shift and cleavage of the peptide backbone[38]. While the two fragments retain the overall structure of mEOS, they are unable to reassemble after mechanical unfolding, resulting in a loss of fluorescence **(Figure 1A)**.

**Figure 1:**
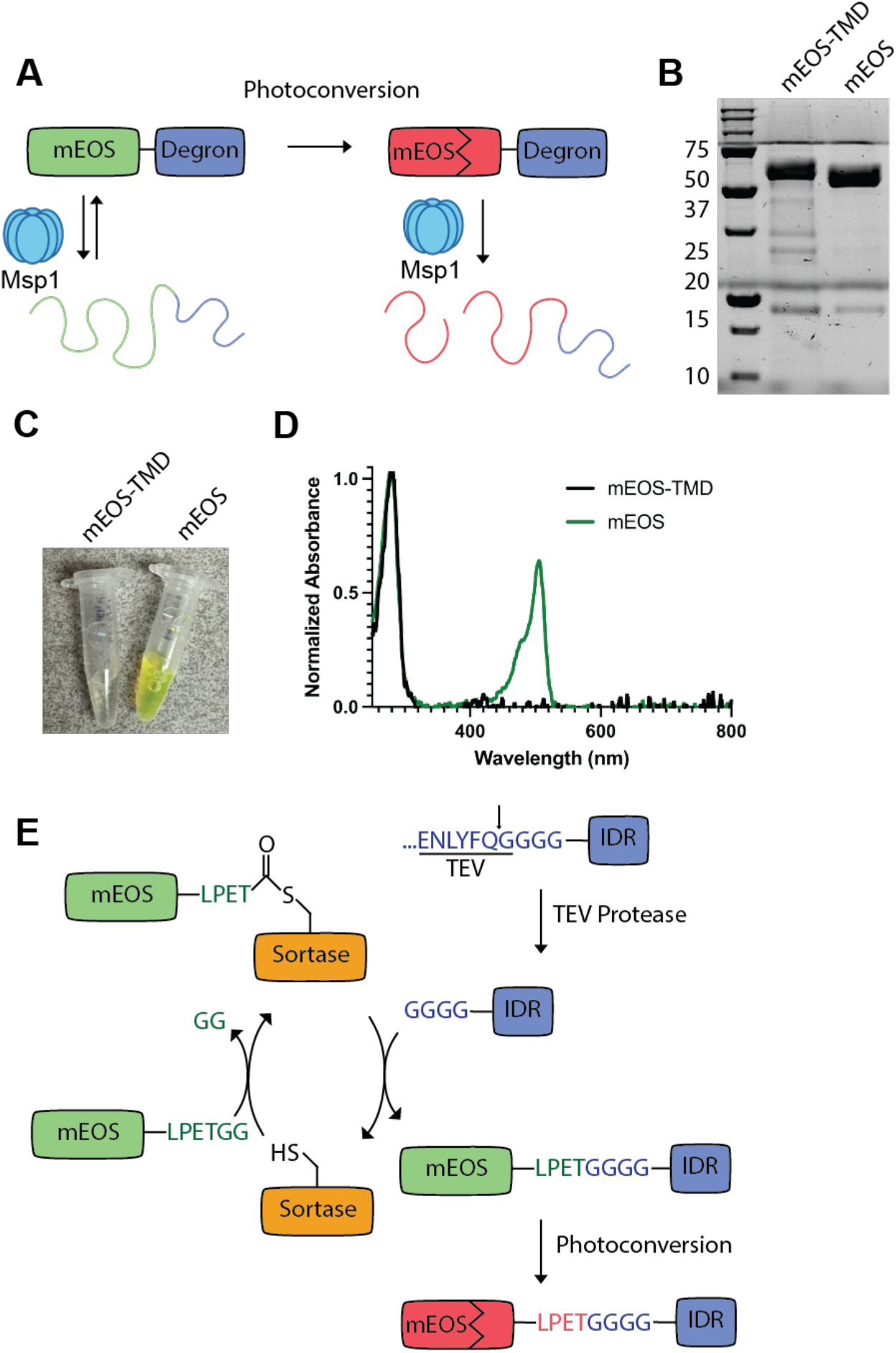
Overview of strategy for fusing Msp1 recognition sequence to mEOS. A) Msp1-mediated unfolding of green (non-photoconverted) mEOS is reversible, making it a poor reporter of Msp1 unfoldase activity. Degron refers to an Msp1 recognition sequence. Photoconversion of mEOS results in a green to red color shift and cleavage of the peptide backbone. Msp1-mediated unfolding of photoconverted mEOS is irreversible. B) SDS PAGE analysis of Ni-NTA purified His_6_-3xFlag-Sumo-mEOS-Sec22TMD-Opsin (mEOS-TMD) and His_6_-3xFlag-Sumo-mEOS-Opsin (mEOS). C) Picture of purified proteins from B. The mEOS-TMD construct has no visible fluorescence. D) Absorbance spectrum of mEOS-TMD and mEOS from B. Absorbance is normalized to 280 nm. E) Overview of strategy for fusing the IDR degron onto mEOS. The IDR degron has a TEV protease site, which is marked by the arrow. TEV cleavage reveals an N-terminal polyglycine sequence. The enzyme Sortase facilitates formation of a peptide bond between the C-terminal LPETGG sequence on mEOS and the N-terminal polyglycine sequence on the IDR degron. Following the sortase reaction, mEOS is photoconverted.

Recent work has demonstrated that soluble mEOS fused to an intrinsically disordered region (mEOS-IDR) can serve as a reporter for Msp1 unfoldase activity[36]. While this assay represents a breakthrough, a limitation is that all validated Msp1 substrates are membrane proteins[1,22,23], whereas mEOS-IDR is a fully soluble protein. Genetically fusing mEOS to a hydrophobic sequence results in a loss of fluorescence, rendering it unusable as an unfolding reporter. An alternative strategy is to separately purify mEOS and an Msp1 recognition sequence and then covalently link these two polypeptides using the enzyme Sortase. Here, we show proof of principle for this strategy.

## Results

### Overview of strategy for covalently attaching IDR degron to mEOS

To adapt mEOS for the study of membrane protein extraction, we first generated a genetic fusion between mEOS and the previously established Msp1 model substrate Sumo-Sec22TMD[27,32]. The full-length construct, His_6_-3xFlag-Sumo-mEOS-Sec22TMD-Opsin, is referred to as mEOS-TMD for simplicity. While the protein was successfully purified with high yield **(Figure 1B)**, we observed no green color **(Figure 1C & D)**. We hypothesized that genetic fusion of mEOS to a hydrophobic sequence disrupts mEOS fluorescence.

To test this hypothesis, we generated an identical construct lacking the hydrophobic transmembrane domain (TMD), His_6_-3xFlag-Sumo-mEOS-Opsin, which we will refer to as mEOS. To control for the possibility that the detergent used in the purification of mEOS-TMD disrupted fluorescence, we purified mEOS using identical buffers and detergents as mEOS-TMD. Purification of this construct resulted in a robust green color, even when purified in the presence of detergent **(Figure 1C & D)**. While the mechanistic details of how the hydrophobic TMD disrupts mEOS fluorescence require further investigation, these results indicate that a direct genetic fusion of mEOS to hydrophobic sequences is not a viable option for producing Msp1 model substrates.

To circumvent this difficulty, we sought to separately purify mEOS and an Msp1 recognition sequence followed by covalent linkage using the enzyme Sortase. Previous work demonstrated that genetically fusing mEOS to an intrinsically disordered region, hereafter referred to as the IDR degron, allows Msp1 to recognize and unfold mEOS[36]. We therefore sought to demonstrate proof of principle for our strategy by separately purifying mEOS and the IDR degron and then covalently linking the two constructs together via Sortase.

Sortase catalyzes the formation of a peptide bond between polypeptides containing a C-terminal LPXTG sequence and an N-terminal polyglycine sequence[39,40]. Our strategy involves purifying mEOS with a genetically encoded C-terminal LPETGG sequence, the optimal sequence for the highly active Sortase pentamutant[41], and generating the IDR degron with an N-terminal polyglycine sequence. As protein translation starts with methionine, we used the TEV protease to expose an N-terminal polyglycine sequence. We chose the TEV protease because it has a recognition sequence of ENLYFQG, with cleavage occurring between glutamine and glycine, thereby leaving the resulting protein with an N-terminal glycine[42,43] **(Figure 1E)**. Sortase then facilitates peptide bond formation between the C-terminal LPETGG sequence on mEOS and the polyglycine sequence on the IDR degron. Finally, the mEOS in the new substrate is photoconverted, resulting in the green to red color shift.

### Purification of mEOS and IDR degron

The IDR degron, His_6_-MBP-TEV-GGGG-IDR-3C-His_6_, was generated by gene synthesis and cloned into a pET28b vector, expressed in *E. coli*, and first purified by Ni-NTA chromatography. The protein was then incubated with TEV protease overnight to remove the His_6_-MBP tag and expose the N-terminal polyglycine sequence for the Sortase reaction. Efficient TEV protease cleavage required the use of a high efficiency TEV protease[43], a 36 h incubation, and the addition of fresh TEV protease after 20 h of incubation.

Following TEV cleavage, we attempted to separate the desired IDR degron from the MBP contaminants, which included cleaved His_6_-MBP, uncleaved substrate (MBP-IDR), and MBP-TEV protease **(Figure 2A)**. We first incubated the sample with amylose resin and collected the flow through. This removed the majority of MBP contaminants.

**Figure 2:**
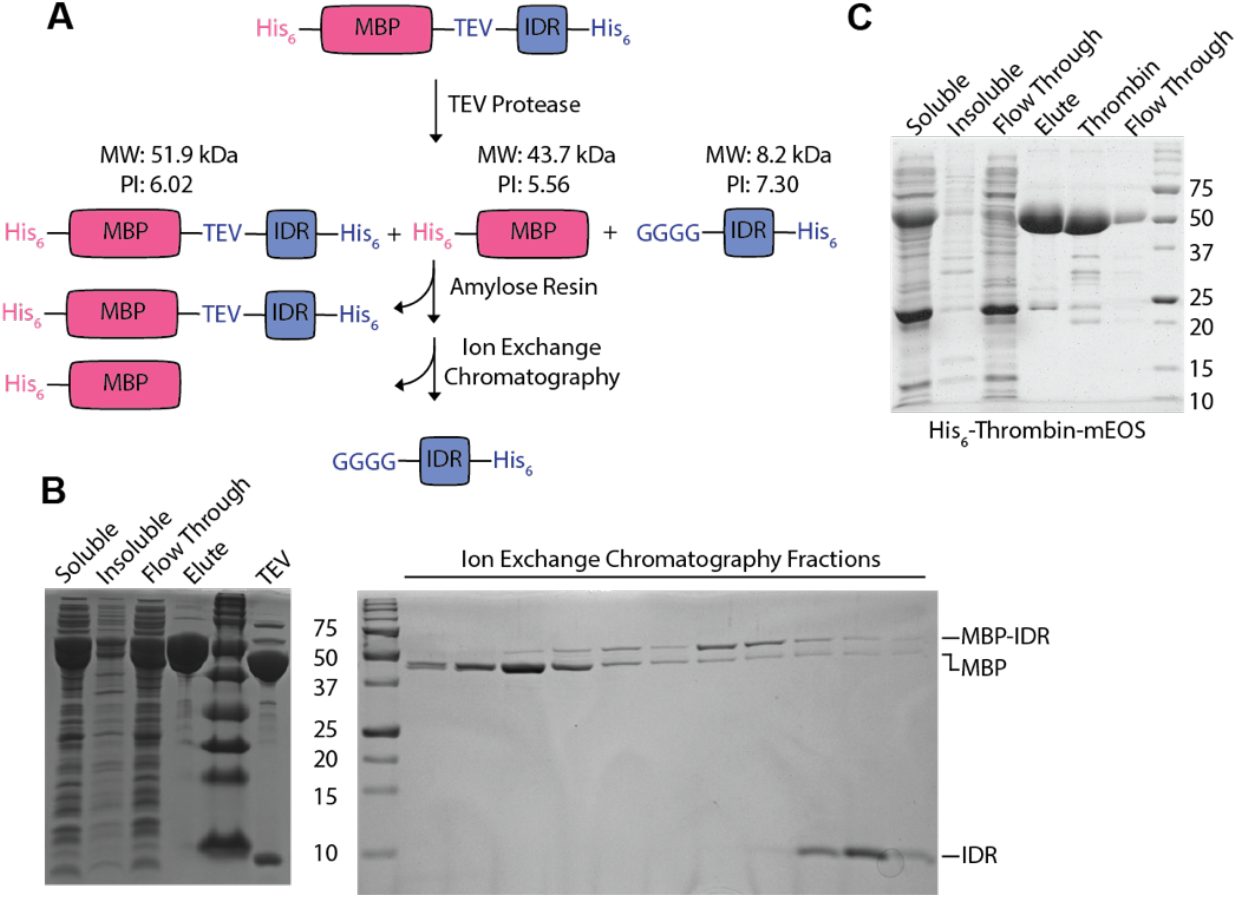
Purification of mEOS and the IDR degron. A) Overview of strategy to purify IDR degron. The IDR degron is initially purified as an N-terminal His_6_-MBP fusion. Following TEV protease cleavage, the IDR degron is separated from the cleaved MBP and uncleaved substrate by passing over amylose resin and then ion exchange chromatography. B) SDS PAGE analysis of IDR degron purification. C) SDS PAGE analysis of mEOS purification.

We attempted to further purify the cleaved substrate from the MBP contaminants by ion exchange chromatography. The cleaved substrate has a predicted isoelectric point (PI) of 7.30, whereas the full-length substrate and cleaved His_6_-MBP have PIs of 6.02 and 5.56 respectively. To facilitate the separation by ion exchange, the Ni-NTA purified substrate was dialyzed into a low salt buffer (Bis-Tris pH 6.0, 25 mM KCl) during TEV protease cleavage. As a result of these two purification steps, we were able to remove the majority of His_6_-MBP and uncleaved substrate (MBP-IDR) **(Figure 2B)**.

The mEOS-LPETGG construct, His_6_-thrombin-mEOS-LPETGG, was generated by gene synthesis and cloned into a pET28b vector. Following expression in *E. coli*, it was purified by Ni-NTA chromatography, cleaved with thrombin overnight, and further purified by size exclusion chromatography. Protein purity, yield, and thrombin cleavage efficiency were all high **(Figure 2C)**.

### Sortase reaction and purification of mEOS-IDR

We decided to use a 3:1 ratio of mEOS to IDR degron to maximize the conversion of the limiting reagent, the IDR degron, into the desired product. We then used the His_6_-tag on the IDR degron to separate the desired reaction product from the unreacted mEOS **(Figure 3A)**. Finally, we performed size exclusion chromatography to separate the desired reaction product from any unreacted IDR degron and the Sortase enzyme. This resulted in relatively pure product with the correct molecular weight **(Figure 3B)**.

**Figure 3:**
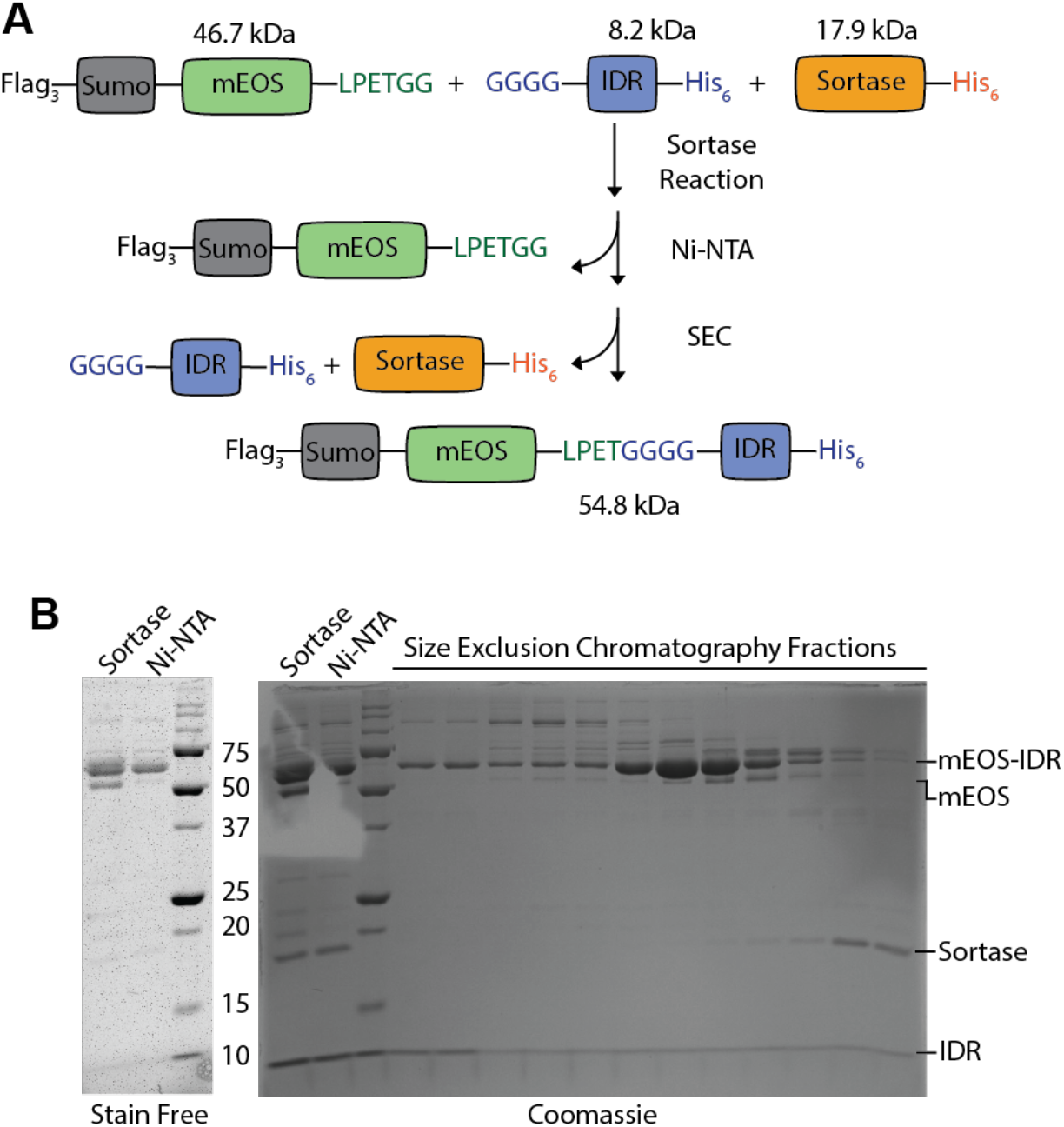
Sortase reaction and purification of mEOS-IDR. A) Diagram of the sortase reaction and purification strategy. Following the sortase reaction, the full-length mEOS-IDR is separated from unreacted mEOS via Ni-NTA chromatography. Size exclusion chromatography is then used to remove Sortase and unreacted IDR degron. B) SDS PAGE analysis of samples following sortase labeling, Ni-NTA enrichment, and size exclusion chromatography. The stain free image on the left is the same gel as on the right, but was aquired before coomassie staining. The IDR lacks tryptophan residues and is therefore not visible on the stain free gel.

### Optimization of mEOS photoconversion

With the pure mEOS-IDR substrate in hand, we next proceeded to photoconvert mEOS. To facilitate photoconversion of milliliter volumes of purified mEOS-IDR, we custom built a photoconversion chamber **(Figure 4A)**. The light source is a 50 W, LED lamp chip with peak emission at 395 nm. A heat sink onto top of the light prevents overheating of the light source. The light is focused through a convex lens with a 5 cm focal length. The lamp and lens are housed in a wooden chamber that protects users from the ultraviolet light. The mEOS-IDR substrate is then placed in a 29 mm glass bottom dish 5 cm from the lens. The glass dish is placed in an ice bucket to prevent sample overheating.

**Figure 4:**
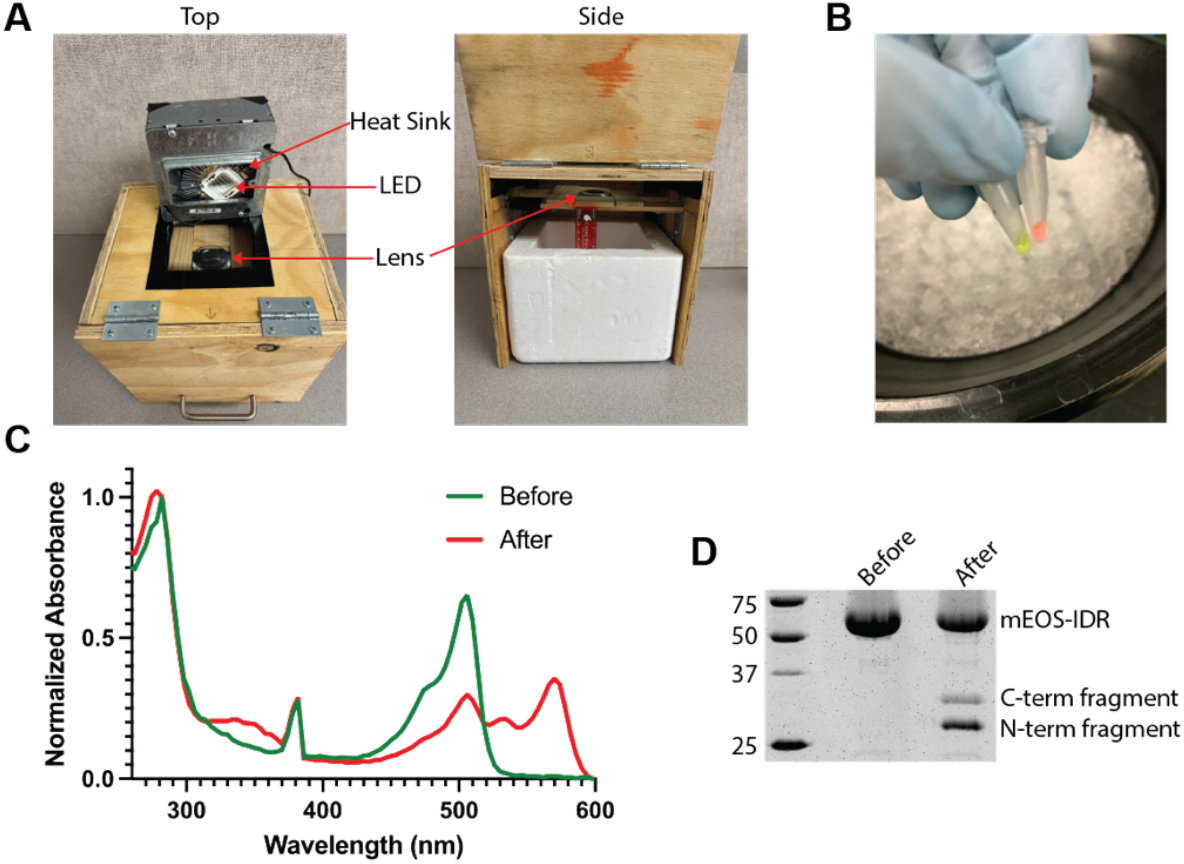
Photoconversion of sortase labeled mEOS. **A)** Pictures of the photoconversion chamber. In the top view, the heat sink and LED are flipped on the side for easy visualization. In the side view, a ruler shows that the lens is 5 cm above the top of the ice bucket, which corresponds to the focal length of the lens. **B)** Picture of sortase labeled mEOS before (left) and after (right) photoconversion shows the green to red color switch. **C)** Absorbance spectrum of mEOS-IDR before and after photoconversion. Absorbance is normalized to 280 nm. **D)** SDS PAGE analysis of sortase labeled mEOS shows cleavage of the peptide backbone in the photoconverted product.

After 30 minutes of exposure to light, we observed a green to red color shift in the mEOS-IDR substrate **(Figure 4B)**. Successful photoconversion was confirmed by absorbance spectra **(Figure 4C)**. Photoconversion causes peptide backbone cleavage in the HYG motif in mEOS, resulting in an N-terminal fragment of 26.8 kDa and a C-terminal fragment of 27.9 kDa. SDS PAGE analysis shows the formation of two bands with the expected molecular weight of the N and C-terminal fragments. Based on band intensity, approximately 10%-20% of the substrate was successfully photoconverted **(Figure 4D)**, which is comparable to previous results with the mEOS-IDR substrate[36].

## Discussion

The photoconvertible fluorescent protein mEOS is a potentially valuable tool for studying substrate unfolding by AAA+ proteins. However, genetically fusing mEOS to a transmembrane domain resulted in a complete loss of fluorescence. As many AAA+ proteins recognize hydrophobic sequences, this severely limits the usability of mEOS as a model substrate for AAA+ unfoldases[44–48]. Here, we describe a strategy that uses Sortase to covalently link mEOS and the IDR degron that were separately purified. We then show that the resulting mEOS-IDR substrate can be successfully photoconverted. Future work will focus on adapting this strategy for use with membrane proteins.

Although proof of principle has been established, the overall substrate design and engineering process could benefit from further optimization to increase yield. The limiting reagent in our reaction was the IDR degron. An alternative strategy is to use an orthogonal tag on the IDR degron rather than relying on a subtractive step such as amylose resin, which could reduce yield via non-specific interactions of the IDR degron with the amylose resin. While the His_6_ tag on the IDR degron facilitated purification after the Sortase reaction, we could not use it for separation from the MBP contaminants due to the N-terminal His_6_ tag on MBP. Future construct optimization of the IDR degron could benefit from replacing the C-terminal His_6_ with small orthogonal affinity tags, such as ALFA-tag of twin-Strep tag [49].

Another potential change is to use a scarless protease, such as NedP1, instead of TEV, which required 36 h of incubation to for 90% cleavage efficiency. The advent of new high efficiency *E. coli* cloning vectors with multiple different tags[50] and the ever-decreasing cost of gene synthesis should facilitate rapid screening of constructs and tags.

A final potential area of optimization is the photoconversion process. The intense light required for photoconversion has the risk of heating the samples, leading to protein unfolding and aggregation. While we mitigated this risk by keeping the samples on ice throughout the photoconversion process, hydrophobic degrons are more aggregation prone and therefore potentially more sensitive to transient overheating. An alternative strategy is to perform the photoconversion before the Sortase reaction.

In conclusion, we present proof of principle for a strategy to covalently attach degrons onto the unfoldase reporter mEOS. We envision this approach could be broadly applicable for other membrane associated AAA+ proteins, offering a powerful tool for advancing biochemical studies on this essential class of enzymes.

## Methods

### Expression and Purification of mEOS constructs

All mEOS constructs were expressed in *E. coli* BL21 DE3 cells with pRIL plasmid in Terrific Broth. Cultures were grown at 37°C until an OD_600_ of 0.6, induced with 0.25 mM isopropyl β-d-1-thiogalactopyranoside (IPTG) for 16 h at 18°C. Cells were harvested by centrifugation and resuspended in buffer supplemented with 0.05 mg/mL lysozyme (Sigma) and 500 U of universal nuclease (Pierce) and lysed by sonication.

For His_6_-thrombin-3xFlag-sumo-mEOS-3C-LPETGG (pArch_252), the cell pellet was resuspended in mEOS Lysis Buffer (25 mM Hepes pH 7.5, 100 mM NaCl, 100 mM KCl, 10 mM MgCl_2_, 20 mM imidazole, 10% glycerol, 0.5 mM EDTA). Following sonication, supernatant was isolated by centrifugation for 30 minutes at 4° C at 18,500 x g and purified by Ni-NTA affinity chromatography (Pierce) on a gravity column. Ni-NTA resin was washed with 30 column volumes (CV) of mEOS Lysis Buffer and eluted with 3 CV mEOS Elution Buffer (25 mM Hepes pH 7.5, 100 mM NaCl, 100 mM KCl, 10 mM MgCl_2_, 250 mM imidazole, 10% glycerol, 0.5 mM EDTA) followed by concentration using a 30 kDa spin concentrator.

For thrombin cleavage, the protein was dialyzed into mEOS Lysis Buffer and thrombin was added at 1:100 ratio and incubated at 4° C for 24 h. The sample was passed over 1 mL of Ni-NTA equilibrated in mEOS Lysis Buffer and the flow through was collected, spin concentrated, and loaded onto a Superdex 200 Increase 10/300 GL column equilibrated in mEOS FPLC Buffer (50 mM Hepes pH 7.5, 50 mM NaCl, 50 mM KCl, 10 mM MgCl_2_, 100 mM imidazole, 5% glycerol, 0.5 mM EDTA). Pure fractions were pooled, spin concentrated, aliquoted, flash frozen in liquid nitrogen, and stored at -80°C.

For the His_6_-thrombin-3xFlag-Sumo-mEOS-Sec22TMD-Opsin (p454_MLW) and His_6_-thrombin-3xFlag-Sumo-mEOS-Opsin (p460_MLW) constructs, the cell pellet was resuspend in mEOS-TMD Lysis Buffer (20 mM Tris pH 7.5, 100 mM NaCl, 20 mM Imidazole, 0.01 mM EDTA, 1 mM DTT). Following sonication, the detergent Lauryldimethylamine-N-oxide (LDAO) was added to a final concentration of 1% and rocked at 4° C for 90 minutes. Supernatant was isolated by centrifugation for 30 minutes at 4° C at 18,500 x g and purified by Ni-NTA affinity chromatography (Pierce) on a gravity column. Ni-NTA resin was washed with 30 CV of mEOS-TMD Lysis Buffer supplemented with 0.1% LDAO. Proteins were eluted with 3 CV mEOS-TMD Elution Buffer (25 mM Hepes pH 7.5, 100 mM NaCl, 100 mM KCl, 10 mM MgCl_2_, 250 mM imidazole, 10% glycerol, 0.5 mM EDTA) supplemented with 0.1% LDAO.

### Expression and Purification of Sortase

Sortase A pentamutant-His_6_ (pArch_066) was expressed in E. coli LOBSTR cells in Terrific Broth. Cultures were grown, shaking at 37°C until OD_600_ of 0.6. Cells were induced with 0.25 mM isopropyl β-d-1-thiogalactopyranoside (IPTG) for 3 h at 30°C. Cells were harvested by centrifugation followed by resuspension in Sortase Lysis Buffer (50 mM Hepes pH 7.5, 300 mM NaCl, 1 mM MgCl_2_) supplemented with 0.05 mg/mL lysozyme (Sigma) and 500 U of universal nuclease (Pierce) and lysed by sonication.

Following sonication, supernatant was isolated by centrifugation for 30 minutes at 4° C at 18,500 x g and purified by Ni-NTA affinity chromatography (Pierce) on a gravity column. Ni-NTA resin was washed with 20 CV of Sortase Lysis Buffer followed by 10 CV of Sortase Wash Buffer (50 mM Hepes pH 7.5, 300 NaCl, 1 mM MgCl_2_, 40 mM imidazole), and eluted with 2 CV of Sortase Elution Buffer (50 mM Hepes pH 7.5, 300 mM NaCl, 1 mM MgCl_2_, 250 mM imidazole). The sample was concentrated by spin concentrator and loaded onto a Superdex200 Increase 10/300GL equilibrated in Sortase FPLC buffer (25 mM Hepes pH 7.5, 150 mM NaCl) for size exclusion. Fractions were analyzed on SDS PAGE, concentrated, aliquoted, flash frozen, and stored at -80° C.

### Expression and MBP-His_7_-superTEV

MBP-His7-superTEV (pArch_066) was expressed in E. coli LOBSTR cells in Terrific Broth. Cultures were grown, shaking at 37°C until OD_600_ of 0.6. Once grown, cells were induced with 0.25 mM isopropyl β-d-1-thiogalactopyranoside (IPTG) for 16 h at 16° C. Cells were harvested by centrifugation followed by resuspension in TEV Lysis Buffer (20 mM Tris pH 7.5, 200 mM NaCl, 1 mM EDTA) supplemented with 0.05 mg/mL lysozyme (Sigma) and 500 U of universal nuclease (Pierce) and lysed by sonication.

Following sonication, supernatant was isolated by centrifugation for 30 minutes at 4° C at 18,500 x g and purified by affinity chromatography with Amylose resin. The Amylose resin was washed with 15 CV of TEV Lysis Buffer and eluted with 2 CV of TEV Lysis Buffer supplemented with freshly added maltose at a final concentration of 10 mM. Sample was concentrated in a spin concentrator and loaded onto Superdex200 Increase 10/300GL equilibrated in TEV FPLC Buffer (20 mM Tris pH 7.5, 100 mM NaCl, 0.1 mM TCEP). Fractions were analyzed on SDS PAGE, concentrated, aliquoted, flash frozen, and stored at -80° C.

### Expression and Purification of IDR degron

The construct His_6_-MBP-TEV-IDR-3C-His_6_ (pArch150) was expressed in *E. coli* BL21 DE3 cells with pRIL plasmid in Terrific Broth. Cultures were grown at 37°C until an OD_600_ of 0.6, induced with 0.25 mM isopropyl β-d-1-thiogalactopyranoside (IPTG) for 16 h at 21°C. Cells were pelleted by centrifugation and resuspended in IDR Lysis Buffer (25 mM Hepes pH 7.5, 150 mM KCl, 1 mM MgCl_2_, 20 mM imidazole, 0.1 mM EDTA, 1 mM DTT) supplemented with 0.05 mg/mL lysozyme (Sigma) and 500 U of universal nuclease (Pierce) and lysed by sonication.

Following sonication, supernatant was isolated by centrifugation for 30 minutes at 4° C at 18,500 x g and purified by Ni-NTA affinity chromatography (Pierce) on a gravity column. Ni-NTA resin was washed with 30 CV of IDR Lysis Buffer, 8 CV of IDR Wash Buffer (25 mM Hepes pH 7.5, 150 mM KCl, 1 mM MgCl_2_, 40 mM imidazole, 0.01 mM EDTA, 0.5 mM DTT), and then eluted with 2 CV IDR Elution Buffer (25 mM Hepes pH 7.5, 150 mM KCl, 1 mM MgCl_2_, 250 mM imidazole, 0.01 mM EDTA).

The protein was dialyzed into Low Salt Buffer (25 mM Bis-Tris pH 6.0, 25 mM KCl, 1mM DTT) and cleaved with MBP-His_7_-superTEV protease (pArch_066) at 1:50 for 36 h at 4°C, refreshing the protease at 20 h. Following TEV cleavage, the sample was passed over 1 mL of packed Amylose resin to remove MBP-contaminants. Flow through was loaded onto an Enrich Q 5×50 anion exchange column (Bio-Rad) equilibrated in Low Salt Buffer. The sample was then washed and eluted with High Salt Buffer (25 mM Bis-Tris pH 6.0, 750 mM KCl, 1mM DTT) using the following wash protocol. Step 1: Linear gradient of 0% to 10% High Salt Buffer over 1 CV. Step 2: Step gradient with 20% High Salt Buffer for 1 CV. Step 3: Step gradient with 100% High Salt Buffer for 1 CV. Pure fractions were pooled and dialyzed into IDR FPLC buffer (50 mM Hepes pH 7.5, 100 mM KCl, 1 mM MgCl_2_, 1 mM DTT, 0.01 mM EDTA, 20 mM imidazole) for 16 h. PEG 20,000 was added to a final concentration of 10% in the dialysis buffer to help concentrate the protein. Concentration was determined via Bradford assay using bovine serum albumin (BSA) for a standard curve. Aliquots were flash frozen in liquid nitrogen and stored at -80° C.

### Sortase Labeling

The sortase reaction contained 5 µM sortase, 45 µM mEOS-LPETGG, and 15 µM of GGGG-IDR. The reaction was brought up to a final volume of 2 mL by addition of Sortase Reaction Buffer (50 mM Hepes pH 7.5, 300 mM KCl, 1 mM MgCl_2_, 0.5 mM TCEP, 10 mM CaCl_2_). The reaction was incubated for 2 hours in a thermomixer at 25° C with 500 rpm.

Following the sortase reaction, the sample was then passed over Ni-NTA resin equilibrated in mEOS Lysis Buffer (25 mM Hepes pH 7.5, 100 mM NaCl, 100 mM KCl, 10 mM MgCl_2_, 20 mM imidazole, 10% glycerol, 0.5 mM EDTA). The resin was washed with 20 CV of mEOS Lysis Buffer and eluted with 2.5 CV of mEOS Elution Buffer (25 mM Hepes pH 7.5, 100 mM NaCl, 100 mM KCl, 10 mM MgCl_2_, 250 mM imidazole, 10% glycerol, 0.5 mM EDTA). The sample was loaded onto a Superdex200 Increase 10/300GL equilibrated in mEOS FPLC Buffer (50 mM Hepes pH 7.5, 50 mM NaCl, 50 mM KCl, 10 mM MgCl_2_, 100 mM imidazole, 5% glycerol, 0.5 mM EDTA). Pure fractions were pool, concentrated by spin concentrator, aliquoted, flash frozen in liquid nitrogen, and stored at -80° C.

### Photoconversion

For photoconversion, 145 μL of mEOS-IDR was placed in a 29 mm glass dish. The dish was placed in an ice bucket 5 cm below the convex lens in the custom built photoconversion chamber. The protein was exposed to light from a 50 W LED lamp chip with a peak emission at 395 nm for 30 minutes. After photoconversion samples were analyzed by absorbance spectrum and SDS PAGE.

**Table 1:** List of plasmids used in this study Glossary.

| Plasmid | Description | Reference |
| --- | --- | --- |
| pArch_150 | His<sub>6</sub>-MBP-TEV-IDR-3C-His<sub>6</sub> | This study |
| pArch_252 | His<sub>6</sub>-thrombin-3xFlag-sumo-mEOS-3C-LPETGG | This study |
| pArch_254 | Sortase A pentamutant-His<sub>6</sub> | Chen et al. 2011[41]<br>Addgene #75144 |
| pArch_066 | MBP-His<sub>7</sub>-superTEV | Keeble et al.<br>2022[43]<br>Addgene #171782 |
| p454_MLW | His<sub>6</sub>-thrombin-3xFlag-sumo-mEOS-Sec22TMD-Opsin tag | This study |
| p460_MLW | His<sub>6</sub>-thrombin-3xFlag-sumo-mEOS-Opsin tag | This study |

## Glossary

AAA+: ATPases Associated with diverse cellular Activities
ATP: Adenosine Triphosphate
GFP: Green Fluorescent Protein IDR: Intrinsically Disordered Region
MBP: Maltose Binding Protein
OMM: Outer Mitochondrial Membrane
TEV: Tobacco Etch Virus
TMD: Transmembrane Domain
TOM: Translocase of the Outer Membrane

## Acknowledgements

The authors wish to thank members of the Wohlever lab for discussions and feedback on the project.

## Author Contributions

Conceptualization: IRW, BAS, DC, AM, MLW; Data curation: IRW, BAS, MLW; Formal analysis: IRW, BAS, MLW; Funding acquisition: MLW; Investigation: IRW, BAS; Methodology: IRW, BAS; Project administration: MLW; Resources: IRW, BAS, MLW; Software: Not applicable; Supervision: AM, MLW; Validation: IRW, BAS; Visualization: IRW, MLW; Writing – original draft: IRW; Writing – review & editing: IRW, BAS, DC, AM, MLW;

